# Epigenetic Regulators Modulate Muscle Damage in Duchenne Muscular Dystrophy Model

**DOI:** 10.1101/211870

**Authors:** Fernanda Bajanca, Laurence Vandel

## Abstract

Histone acetyl transferases (HATs) and histone deacetylases (HDAC) control transcription during myogenesis. HDACs promote chromatin condensation, inhibiting gene transcription in muscle progenitor cells until myoblast differentiation is triggered and HDACs are released. HATs, namely CBP/p300, activate myogenic regulatory and elongation factors promoting myogenesis. HDAC inhibitors are known to improve regeneration in dystrophic muscles through follistatin upregulation. However, the potential of directly modulating HATs remains unexplored. We tested this possibility in a well-known zebrafish model of Duchenne muscular dystrophy. Interestingly, CBP/p300 transcripts were found downregulated in the absence of Dystrophin. While investigating CBP rescuing potential we observed that *dystrophin*-null embryos overexpressing CBP actually never show significant muscle damage, even before a first regeneration cycle could occur. We found that the pan-HDAC inhibitor trichostatin A (TSA) also prevents early muscle damage, however the single HAT CBP is as efficient even in low doses. The HAT domain of CBP is required for its full rescuing ability. Importantly, both CBP and TSA prevent early muscle damage without restoring endogenous CBP/p300 neither increasing follistatin transcripts. This suggests a new mechanism of action of epigenetic regulators protecting *dystrophin*-null muscle fibres from detaching, independent from the known improvement of regeneration upon damage of HDACs inhibitors. This study builds supporting evidence that epigenetic modulators may play a role in determining the severity of muscle dystrophy, controlling the ability to resist muscle damage. Determining the mode of action leading to muscle protection can potentially lead to new treatment options for muscular dystrophies in the future.

## INTRODUCTION

There is no cure to date for muscular dystrophies caused by absent or malfunctioning Dystrophin, be it lethal Duchenne or milder Becker type. Current therapies aim at alleviating the symptoms and delaying the progression of a disease that can be life threatening. One promising pharmacological treatment is to inhibit histone deacetylases (HDACs) (Consalvi et al., 2011). HDACs can regulate gene transcription in muscle progenitor cells, by controlling the activity of myogenic regulatory factors and MEF2 family proteins (McKinsey et al., 2000; Palacios and Puri, 2006). HDACs promote chromatin condensation, inhibiting gene transcription until myoblast differentiation is triggered and HDACs are released. Similarly, blocking HDACs leads to hyper acetylated chromatin, inhibiting condensation and therefore facilitating transcription. HDAC inhibitors ameliorate the dystrophic phenotype by promoting myogenesis and improving regeneration in dystrophic muscles (Consalvi et al., 2011; Giordani and Puri, 2013; Iezzi et al., 2002). Several studies show that follistatin upregulation by HDAC inhibitors is responsible for boosting regeneration in dystrophic muscles (Iezzi et al., 2004; Mozzetta et al., 2013). While studies have been focusing on blocking HDACs to promote hyper acetylated chromatin and transcription, the potential of directly modulating histone acetyl transferases (HATs) on a muscle dystrophy context remains unexplored. CBP (CREB Binding Protein) is a nuclear transcriptional co-activator with HAT activity, belonging to the p300/CBP family (Bedford et al., 2010). It is ubiquitously expressed and acetylates both histone and non-histone proteins to regulate transcription. CBP was shown to functionally activate MyoD by acetylation and to directly interact with MEF2C (Giordani and Puri, 2013). CBP expression in zebrafish muscle was reported recently (Batut et al., 2015). We used the zebrafish model, extensively characterised and widely used for studying muscular dystrophy (Bajanca et al., 2015), to explore the effects of overexpressing CBP in a Dystrophin-null background. We have found that overexpressing this single HAT rescues the dystrophic phenotype as efficiently as blocking HDACs. Moreover, we observed that both treatments inhibit the early appearance of dystrophic fibres, prior to any effect on regeneration could take place. We report a follistatin-independent mechanism protecting against fibre damage.

## RESULTS

To test whether Dystrophin absence affects HAT's expression we performed quantitative real-time PCR (qPCR) for *p300* and/or *CBP* (*crebbp*) transcripts (Figure 1A). Two-way ANOVA shows highly significant differences [F(1,24)=31.5, p<0.0001] between HAT genes expression in controls and embryos injected with well described Dystrophin morpholino cocktail (also see Supplementary Methods, Supplementary Figure 1A). Multiple comparisons indicate strong downregulation of *crebbpb* and *p300b* and milder but significant decrease of *p300a* expression (*p* values: *crebbpb* = 0.0010, *p300a* = 0.0247, *p300b* = 0.0045). Therefore, the absence of Dystrophin significantly affects HAT expression.

**Figure 1.**
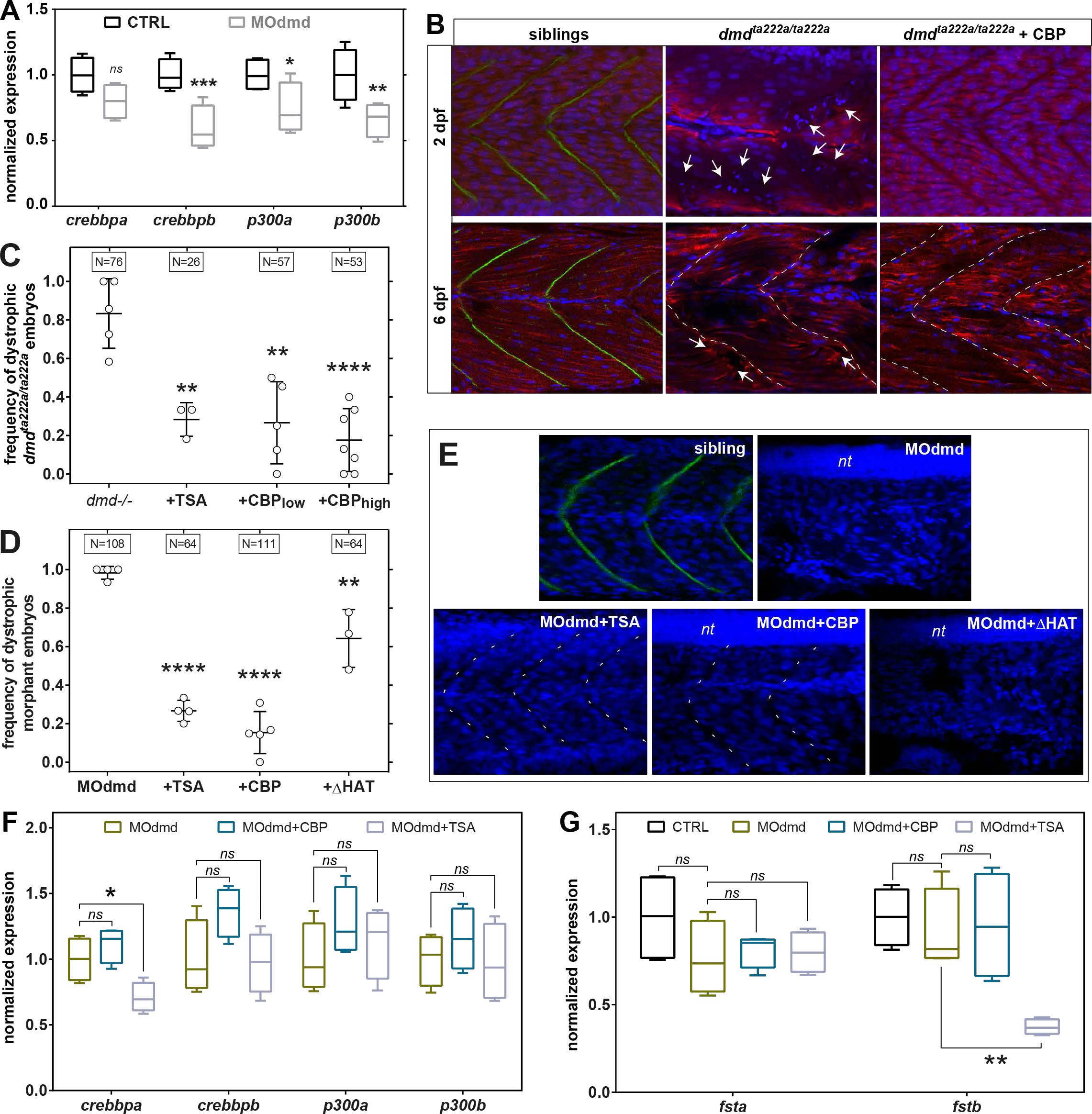
Epigenetic regulators in zebrafish dystrophic embryos. **(A)** Quantitative PCR analysis of p300/CBP transcript expression at 2 days post fertilisation (dpf) in control and morpholino-depleted embryos (MOdmd). **(B)** Confocal sections of *Tg(actc1b:mCherry)*^pc4^ embryos immunostained for Dystrophin (green). Well organised pattern of muscle fibres (mCherry, red) and nuclei (DAPI, blue) in siblings, while heavily disrupted in *dmd*^ta222a/ta222a^ embryos (arrows), showing collapsed and wavy fibres which ends misalign with somatic borders (dashed lines). Overexpressing CBP (*dmd*^ta222a/ta222a^ + CBP; injection of 10^7^ RNA molecules of plasmid per embryo at one cell stage) has a clear positive effect on dystrophic embryos both at 2 and 6 dpf. **(C, D)** Frequency of dystrophic phenotype on 2 dpf Dystrophin-*null* embryos, either on *dmd*^ta222a/ta222a^ mutants (C) or depleted with MOdmd (D). Overexpressing CBP (low = 10^7^, high = 10^8^ molecules of plasmid RNA injected per embryo) rescues the dystrophic phenotype of both *dmd*^ta222a/ta222a^ (C) and morphant (D; 10^7^ molecules) embryos as efficiently as blocking HDACs (TSA), even at lower concentrations (C,D). A mutated CBP form lacking the HAT domain (ΔHAT; 10^7^ molecules) significantly decreases the rescue efficiency (C). Individual values for independent experiments are plotted, bars represent the average ± SD and boxes above plots indicate total number of embryos analysed. **(E)** Representative confocal sections of 2 dpf embryos immunostained for Dystrophin (green) to confirm Dystrophin presence in siblings and depletion in MOdmd embryos, counterstained with DAPI (blue). Both TSA (200 nM) and CBP (10^7^ molecules) are able to rescue the muscle phenotype of MOdmd embryos, while ΔHAT injected embryos most often show muscle damage (10^7^ molecules). *nt* = neural tube. **(F,G)** Quantitative PCR analysis of *crebbp*, *p300* and *follistatin* endogenous transcripts expression at 2 dpf. See text for details on ANOVA (A,F,G) or *T* tests (B,C), performed with cut-offs: (*) < 0.05, (**) < 0.005, (***) < 0.001, (****) < 0.0001. *ns* = non significant.

Taking these results, we set to determine whether the dystrophic phenotype could be rescued by overexpressing RFP-tagged murine CBP in *dystrophin*-null embryos (*dmd^ta222a/ta222a^*). Nuclear and dose dependent expression of the transgene was confirmed (Supplementary Figure 1B,C). To test the hypothesis that CBP overexpression improves regeneration, embryos were analysed at 2 dpf, as fibre tips start detaching from somitic borders (Bajanca et al., 2015), and 6 dpf, when regeneration upon damage is underway (Gurevich et al., 2016; Pipalia et al., 2016). While *dystrophin*-null embryos show characteristic dystrophic muscles from early stages, overexpressing CBP clearly reduced damage not only at later stages but, unexpectedly, also at 2 dpf (Figure 1B). Fast swimming was also restored, a characteristic failure of dystrophic embryos. Absence of damage as early as 2 dpf cannot be explained by stimulating regeneration, which takes several days (Gurevich et al., 2016; Pipalia et al., 2016), suggesting that CBP overexpression may act at an earlier phase preventing muscle degeneration in the first place. We focused on investigating this observation further and testing whether only CBP overexpression or also HDAC inhibition could have a protective effect. Different concentrations of CBP were tested and rescue at 2 dpf compared with TSA treatment, previously used in zebrafish and considered the most efficient pan-HDAC inhibitor (Consalvi et al., 2011; Johnson et al., 2013) (Figure 1C-E; Supplementary Figure 1D). On average, 83% of *dystrophin*-null (*dmd^ta222a/ta222a^*) embryos showed clear dystrophic muscles at 2 dpf (Figure 1C, *dmd*^-/-^). The prevalence of a dystrophic signature was drastically reduced to 28% upon treatment with TSA and 27% when a low dose of CBP (+CBP_low_) was used. A 10-fold increase of the CBP dose (+CBP_high_) improves the rescue significance (p<0.0001). Therefore, CBP overexpression is at least as efficient as HDACs inhibition in protecting *dystrophin-null* muscles from damage. Similar results were obtained in the Dystrophin morphant background for both HDACs inhibitor and CBP (Figure 1D,E). Immunostaining for RFP-CBP shows that rescue is obtained in morphants even when detectable levels are seen in few muscle nuclei (Supplementary Figure 1D). Overexpression of a CBP form lacking active HAT domain leads to mild rescue indicating that the acetylase activity of CBP is essential for full rescue (+ΔHAT; Figure 1D,E).

Since CBP/p300 expression is generally downregulated in Dystrophin-depleted embryos (Figure 1A), we tested whether CBP overexpression or TSA act though increasing endogenous *crebbp/p300* levels. Dystrophin-depleted embryos were co-injected with CBP (10^7^ molecules) or treated with TSA and qPCR was performed at 2 dpf (Figure 1F; Supplementary Figure 1E). One-way ANOVA revealed statistically significant differences between treatment groups for *crebbpa* expression [F (2, 9) = 8.915, p = 0.0073**], Fisher's LSD post-hoc test finding the decrease upon TSA treatment statistically significant (p = 0.0164*). Independent one-way ANOVAs revealed absence of statistically significant differences in the expression of *crebbpb* [F (2, 9) = 3.340, p = 0.0823], *p300a* [F (2, 9) = 1.099, p = 0.3741] and *p300b* [F (2, 9) = 0.6550, p = 0.5425] transcripts. These results show that neither CBP nor TSA significantly increase any of the transcripts, *crebbpa* being even decreased upon TSA treatment (Figure 1F). Therefore, rescue is not due to restoring endogenous CBP/p300 transcripts levels.

While the mechanism of action of HDAC inhibition leading to improved regeneration is not fully understood, follistatin upregulation is a key event (Iezzi et al., 2004; Mozzetta et al., 2013). We questioned whether the follistatin pathway could also be involved in the early protection against muscle damage observed in TSA treatment and CBP overexpression. Quantitative analysis of the two zebrafish follistatin transcripts, *fsta* and *fstb*, expression was performed at 2 dpf (Figure 1G). Dystrophin depletion does not significantly affect *fsta* or *fstb* levels. There were no statistically significant differences in the expression of *fsta* between groups as determined by one-way ANOVA [F (3, 12) = 1.288, p = 0.3232]. The expression of *fstb* is statistically significantly different between groups [F (3, 12) = 7.680, p = 0.0040**], Fisher's LSD post-hoc test finds the decrease upon TSA treatment statistically significant (p = 0.0013**). Summarising, TSA treatment significantly downregulates the expression of *fstb* while not affecting *fsta* and both transcripts remain unaltered by CBP overexpression. Therefore, the protection against muscle damage granted by CBP overexpression or TSA treatment does not rely on stimulating follistatin transcription but likely on a follistatin-independent pathway.

## FINAL REMARKS

This study shows for the first time that either CBP overexpression or pan-HDAC inhibition are able to prevent, or at least delay, early stages of muscle damage in zebrafish dystrophic muscles. This observation adds up to the recognised positive effect of HDAC inhibitors on muscle regeneration upon damage (Consalvi et al., 2011). Other authors observed that treating mouse dystrophic muscles (mdx) with deacetylase inhibitors was able to confer resistance to contraction-coupled degeneration (Minetti et al., 2006). However, this was suggested to be mediated by follistatin through satellite cell number increase. We show that damage protection occurs independently from regeneration. Follistatin upregulation is not involved in protecting early zebrafish muscle from damage, supporting the idea that the two rescuing mechanisms triggered by HDAC inhibition are independent. Interestingly, overexpression of a single HAT, CBP, achieves a similar level of protection as pan-HDAC inhibition. The strong downregulation of *fstb* by TSA but not CBP should be investigated in future work to determine whether these agents may act through different mechanisms of action. Our results prompt for investigating similar mechanisms in other animal models and identifying new therapeutic targets, possibly with more restricted and controlled effects than HDAC inhibitors. This study strengthen the idea that epigenetics plays a role in the progression of symptoms in some patients with Dystrophin mutations. This could explain the variation in severity in Becker muscular dystrophy patients with identical mutation and the variable progression of DMD. Doubt persists on the mechanism leading to protection. To further understand the mechanisms of action triggered by either HDAC inhibition or CBP a high throughput analysis needs to be employed in future studies.

## MATERIALS AND METHODS

### Animals

Fish used were wild-type Danio rerio, *dmd*^ta222a/+^ and *dmd*^ta222a/+^ crossed to *Tg(actc1b:mCherry)*^pc4^ to facilitate identifying dystrophic muscles. All animals were handled in a facility certified by the French Ministry of Agriculture (approval ID B-31-555-10) and in accordance with the guidelines from the European directive on the protection of animals used for scientific purposes (2010/63/UE), French Decret 2013–118. Staging and rearing was performed as described (Westerfield, 1995). Plasmids and morpholinos were injected into 1-cell stage embryos at the indicated concentrations ((Johnson et al., 2013); see text). Phenolthiourea (0.003%) was added to inhibit pigmentation and facilitate muscle analysis. Dystrophic muscles were characterised by abnormal birefringence, supported by observing disruption of the actin reporter pattern when *Tg(actc1b:mCherry)*^pc4^ fish were used, and impaired movement (Bajanca et al., 2015; Berger et al., 2012). The absence of Dystrophin expression in rescued embryos was confirmed by immunohistochemistry.

### Plasmids, morpholinos and HDACs inhibition

pCS2+mRFP-CBP and pCS2+mRFP-ΔHAT plasmids were created by digesting by BamH1 full length CBP of mouse origin, or CBP deleted for amino acids 1430 to 1570 from pCMV-HA-CBP and pCMV-HA-CBPΔHAT (provided by A. Harel-Bellan) and inserting them into pCS2+mRFP-N1 vector (Addgene) digested by BamH1. Injections were performed at one-cell stage with the indicated amounts of *RFP-CBP* and *RFP-CBPΔHAT* RNA. Mouse CBP shares 85% protein identity to both zebrafish CBP-A (the product of *crebbpb*) and zebrafish CBP-B (the product of *crebbpa*), according to the standard NCBI BLAST tools. Zebrafish protein CREBBPA (also called CREBBPb or CBP-B) corresponds to protein ID ENSDARP00000081684, while CREBBPB protein ID (also called CREBBPa or CBP-A) corresponds to ENSDARP00000086306.

Morpholinos were from Gene-tools and were used as a cocktail as described previously (*dmd*-MO1 at 4 ng per embryo and *dmd*-MO6 at 7.5ng/embryo;(Johnson et al., 2013).

Embryos were exposed to Trichostatin A (TSA, Sigma) for 24 hours, from 24 to 48 hpf. TSA was added to fish water for a final concentration of 200 nM as described previously (He et al., 2014; Johnson et al., 2013).

### Real time quantitative PCR

Four samples were analysed for each condition, each sample consisting of 15 embryos at 2 days post fertilisation (dpf) for a total of 60 embryos per condition. Total RNAs were extracted with the EZNA total RNA kit I (Omega Bio-tek), and reverse-transcribed with the qScript cDNA synthesis kit (Quanta biosciences) according to the supplier’s instructions. A C1000™ Thermal Cycler with CFX96™ Real time System (BioRad) was used to perform the qPCR and analyses were performed on BioRad CFX Manager 2.0, according to the manufacturer’s instructions. SsoFast^™^ EvaGreen^®^ Supermix (Bio-Rad) was used according to the manufacturer’s instructions. The following primer sequences were used: EF1α Fwd 5’-GAT GCA CCA CGA GTC TCT GA -3’; EF1α Rev 5’-TGA TGA CCT GAG CGT TGA AG -3’; m-crebbp Fwd 5'-TGC CAA GTT GCC CAT TGT G -3'; m-crebbp Rev 5'-TTG TTG GTT TCG CTT GTC ACT- 3' ; z-crebbpa Fwd 5’-CGA AAA GTG GAA GGG GAC AT -3’; z-crebbpa Rev 5’-TTC TCT TCC AGC TCT TTC TGG -3’; z-crebbpb Fwd 5’-CAG GTT CCT CAA GGG ATG G -3’; z-crebbpb Rev 5'-CCA TCA TGG CTT GAG CTT G -3’; p300a Fwd 5'-CAC CTT CCT CAA CAC CAC AGT -3’; p300a Rev 5'-GCA TAG CAT TCT GGT CTG CTC -3'; p300b Fwd 5'-ATA TGG CCG TCA GGG TTT ATC -3'; p300b Rev 5'-CTC GTG TCT CCA GAA AGT TGC -3'; fsta Fwd 5’-GAT GCA AAA TGA ACA GGA GGA -3’; fsta Rev 5’-GAC TTC AAA AGG GCA CAT TCA -3’; fstb Fwd 5’-GCA TGG ACT GAG GAG GAT GTA -3’; fstb Rev 5’-CAC CTC TTT CCA GAA CCA CAA -3’. All experiments included a standard curve for each primer pair and control for genomic DNA contamination. Data were normalized to *EF1α* and expressed relative to the respective control that is set as 1 in boxplot with whiskers representing the median, 0.25 percentiles, maximum and minimum values.

### Immunohistochemistry

Standard protocols were used. Embryos were fixed in cold methanol for dystrophin staining, or otherwise in paraformaldehyde 4%. Primary antibodies were mouse anti-Dystrophin (MANDRA1 (7A10), Developmental Studies Hybridoma Bank) and rat anti-RFP (Chromotek). Secondary antibodies were goat anti-mouse Alexa-488 and goat anti-rat Alexa-546 (Molecular Probes). DAPI was used to counterstain nuclei.

### Microscopy, software and statistics

An inverted Zeiss 710 LSM with a 20x/1.0 W Plan-Apochromat and a 40x/1.3 Plan-Apochromat objective were used for *Z*-stack acquisition. Acquisition and maximum intensity projections were made with ZEN 2009/2010 (Zeiss). Images were uniformly contrasted with Adobe Photoshop CS6. Illustrations were made in Adobe Illustrator CS6. GraphPad Prism 6 was used for graph plotting and statistical analyses.

## ACKNOWLEDGEMENTS

We thank Dr. A. Harel-Bellan for plasmids, the Toulouse Rio Imaging (TRI) platform and the fish facility of the CBD. This work was supported by grants from the Fondation de l’Association pour la Recherche contre le Cancer (Fondation ARC); Association Française contre les Myopathies (AFM); Alliance pour la Vie et pour la Santé (AVIESAN) and Institut National du Cancer (INCa) to L.V and a fellowship from the Fondation pour la Recherche Médicale (FRM) to F.B.

## BIBILIOGRAPHY

Bajanca, F., Gonzalez-Perez, V., Gillespie, S. J., Beley, C., Garcia, L., Theveneau, E., Sear, R. P. and Hughes, S. M. (2015). In vivo dynamics of skeletal muscle Dystrophin in zebrafish embryos revealed by improved FRAP analysis. Elife 4,.

Batut, J., Duboé, C. and Vandel, L. (2015). Expression patterns of CREB binding protein (CREBBP) and its methylated species during zebrafish development. Int. J. Dev. Biol. 59, 229–34.

Bedford, D. C., Kasper, L. H., Fukuyama, T. and Brindle, P. K. (2010). Target gene context influences the transcriptional requirement for the KAT3 family of CBP and p300 histone acetyltransferases. Epigenetics 5, 9–15.

Berger, J., Sztal, T. and Currie, P. D. (2012). Quantification of birefringence readily measures the level of muscle damage in zebrafish. Biochem. Biophys. Res. Commun. 423, 785–8.

Consalvi, S., Saccone, V., Giordani, L., Minetti, G., Mozzetta, C. and Puri, P. L. (2011). Histone deacetylase inhibitors in the treatment of muscular dystrophies: epigenetic drugs for genetic diseases. Mol. Med. 17, 457–65.

Giordani, L. and Puri, P. L. (2013). Epigenetic control of skeletal muscle regeneration: Integrating genetic determinants and environmental changes. FEBS J. 280, 4014–4025.

Gurevich, D. B., Nguyen, P. D., Siegel, A. L., Ehrlich, O. V., Sonntag, C., Phan, J. M. N., Berger, S., Ratnayake, D., Hersey, L., Berger, J., et al. (2016). Asymmetric division of clonal muscle stem cells coordinates muscle regeneration in vivo. Science (80-.). 353, aad9969–aad9969.

He, Y., Mei, H., Yu, H., Sun, S., Ni, W. and Li, H. (2014). Role of histone deacetylase activity in the developing lateral line neuromast of zebrafish larvae. Exp. Mol. Med. 46, e94.

Iezzi, S., Cossu, G., Nervi, C., Sartorelli, V. and Puri, P. L. (2002). Stage-specific modulation of skeletal myogenesis by inhibitors of nuclear deacetylases. Proc. Natl. Acad. Sci. U. S. A. 99, 7757–62.

Iezzi, S., Di Padova, M., Serra, C., Caretti, G., Simone, C., Maklan, E., Minetti, G., Zhao, P., Hoffman, E. P., Puri, P. L., et al. (2004). Deacetylase inhibitors increase muscle cell size by promoting myoblast recruitment and fusion through induction of follistatin. Dev. Cell 6, 673–84.

Johnson, N. M., Farr, G. H. and Maves, L. (2013). The HDAC Inhibitor TSA Ameliorates a Zebrafish Model of Duchenne Muscular Dystrophy. PLoS Curr. 5,.

McKinsey, T. A., Zhang, C. L. and Olson, E. N. (2000). Activation of the myocyte enhancer factor-2 transcription factor by calcium/calmodulin-dependent protein kinase-stimulated binding of 14-3-3 to histone deacetylase 5. Proc. Natl. Acad. Sci. 97, 14400–14405.

Minetti, G. C., Colussi, C., Adami, R., Serra, C., Mozzetta, C., Parente, V., Fortuni, S., Straino, S., Sampaolesi, M., Di Padova, M., et al. (2006). Functional and morphological recovery of dystrophic muscles in mice treated with deacetylase inhibitors. Nat. Med. 12, 1147–1150.

Mozzetta, C., Consalvi, S., Saccone, V., Tierney, M., Diamantini, A., Mitchell, K. J., Marazzi, G., Borsellino, G., Battistini, L., Sassoon, D., et al. (2013). Fibroadipogenic progenitors mediate the ability of HDAC inhibitors to promote regeneration in dystrophic muscles of young, but not old Mdx mice. EMBOMol. Med. 5, 626–639.

Palacios, D. and Puri, P. L. (2006). The epigenetic network regulating muscle development and regeneration. J. Cell. Physiol. 207, 1–11.

Pipalia, T. G., Koth, J., Roy, S. D., Hammond, C. L., Kawakami, K. and Hughes, S. M. (2016). Cellular dynamics of regeneration reveals role of two distinct Pax7 stem cell populations in larval zebrafish muscle repair. Dis. Model. Mech. 9, 671–684.

Westerfield, M. (1995). The Zebrafish Book -a guide for the laboratory use of zebrafish (Danio rerio). 3 edn. University of Oregon Press.

